# A Glance into the Destiny of Transcriptomic Activity, Embodied by the HOX Genes, in Neonatal and Aging Dermal Cells

**DOI:** 10.1101/2023.06.28.546883

**Authors:** Doyeong Ko, Minji Kim, Youn-Hwa Nho, Dong-Geol Lee, Seunghyun Kang, Kyudong Han, Seyoung Mun, Misun Kim

## Abstract

Skin is an organ having a crucial role in the protection of muscle, bone, and internal organs and undergoing continuous self-renewal and aged. The growing interest in the prevention of skin aging and rejuvenation has sparked a surge of industrial and research studies focusing on the biological and transcriptional changes that occur during skin development and aging. In this study, we aimed to identify transcriptional differences between two main types of human skin cells: the HDFs and the HEK isolated from 30 neonatal and 30 adults (old) skin. Through differentially expressed gene (DEG) profiling using DEseq2, 604 up-, and 769 down-regulated genes were identified in the old group. The functional classification analysis using Metascape Gene Ontology and Reactome pathway was performed. We report the systematic transcriptomic changes in key biological markers involved in skin formation and maintenance and a unique difference in *HOX* gene families which are important for developing embryonic formation and regulating numerous biological processes. Among the 39 human *HOX* genes, 10 genes (*HOXA10*, *11*, *13*, *HOXB13*, *HOXC11*, and *HOXD9*-*13*) were significantly down-regulated, and 25 genes *HOXA2*-*7*, *HOXB1*-*9*, *HOXC4*-*6* and *8*-*9*, and *HOXD1*,*3*,*4* and *8*) were up-regulated, especially in the old HDFs. We have successfully established a correlation between *HOX* genes and the process of skin aging, thereby proposing *HOX* genes as a novel marker for assessing skin aging. Our findings provide compelling evidence supporting the involvement of *HOX* genes in this biological phenomenon such as skin aging.

## INTRODUCTION

The skin is the body’s largest organ and performs many vital functions. It consists of three primary layers: the outermost epidermis, the middle dermis, and the deepest hypodermis. The epidermis provides a protective anatomical barrier against environmental hazards such as UV radiation, toxins, and microorganisms. It comprises different cell layers, including keratinocytes which synthesize waterproof keratin, and melanocytes which relate to skin pigmentation (Elias and Choi 2005; Proksch et al. 2008; Arda et al. 2014). The dermis, located beneath the epidermis, is a sophisticated layer comprised of various cell types. Among these are fibroblasts, responsible for synthesizing crucial proteins like collagen and elastin that contribute to the skin’s strength and elasticity. Furthermore, the dermis encompasses an intricate network of blood vessels, lymph vessels, and nerves that play essential roles in temperature regulation and provide nutrients and oxygen to the skin. The hypodermis, the deepest layer of skin, consists of adipose and connective tissue and helps to regulate body temperature while providing insulation and cushioning for the body (Arda et al. 2014; Brown and Krishnamurthy 2021). Their harmonious interaction between the dermis and epidermis plays a vital role in maintaining the normal functioning of the skin and preventing damage from external factors. As humans age, changes in skin structure and function accumulate, and the epidermis of aging skin becomes thinner and less efficient in its barrier function. (Ghadially et al. 1995; Waller et al. 2005). Aged skin also has fewer fibroblasts in the dermis, leading to alterations in and degradation of the extracellular matrix (ECM), which manifests as progressed dermal thinning, increased wrinkling, and a loss of elasticity (Shuster et al. 1975; ContetlJAudonneau et al. 1999; Waller et al. 2005; Varani et al. 2006). Additionally, the contact area between the dermis and epidermis decreases and flattens, causing insufficient nutrition supply to the epidermis and reducing the basal cell proliferation capability (Lavker et al. 1989; Varani et al. 2006; Farage et al. 2007; Tobin 2017). Cutaneous aging is accompanied by decreased proliferative ability of skin cells, including keratinocytes, fibroblasts, and melanocytes, which is called cellular senescence (Lavker et al. 1989; Ressler et al. 2006; Victorelli et al. 2019). Skin samples from human donors of different ages show that senescence marker p16INK4a positive cells increase with age in the dermal fibroblasts and epidermal keratinocytes, indicating that aged skin contains more senescent cells (Ressler et al. 2006; Waaijer et al. 2012). These cells contribute to the structural and functional changes in the skin, leading to wrinkles, age spots, and a dull or uneven complexion. As keratinocytes age, they may produce fewer of the proteins and lipids that maintain the skin’s barrier function, leading to dryness and increased susceptibility to damage (Wang and Dreesen 2018; Wang et al. 2020). Senescent fibroblasts in the dermis produce less ECM components, such as collagen and elastin, and more enzymes that break down the extracellular matrix, such as MMPs and elastases, which lead to a loss of skin elasticity and the formation of wrinkles (Waaijer et al. 2016; Wang and Dreesen 2018; Ezure et al. 2019; Lee et al. 2021). While senescence is a natural part of the aging process, environmental factors (e.g., UV radiation, pollution) and lifestyle (e.g., smoking, a poor diet) can accelerate this process by increasing oxidative stress and inflammation in the skin, which can contribute to cellular aging and the breakdown of skin structure and function (Wang and Dreesen 2018; Krutmann et al. 2021; Lee et al. 2021). Several studies have indicated that the accumulation of senescent cells in the skin is linked to age-related changes like decreased elasticity and the development of wrinkles. D.J. Baker demonstrated that cellular senescence is associated with age-related phenotypes and eliminating these cells can delay aging-associated disorders (Baker et al. 2011). In the epidermis, the increase in senescent cells with age is associated with changes in epidermal thickness and facial wrinkles during skin aging (Waaijer et al. 2016; Rübe et al. 2021). The accumulation of senescent fibroblasts in the dermis is also correlated with wrinkle formation and morphological changes in the elastic fibers of the skin (Ressler et al. 2006; Waaijer et al. 2016). Topical treatment with rapamycin has been shown to reduce the number of p16INK4a-positive senescent cells in human skin and decrease wrinkles and increase the integrity of the basement membrane (Chung et al. 2019). Moreover, senescent fibroblasts also contribute to hyperpigmented disorders such as melasma, and the removal of these cells through radiofrequency therapy can increase procollagen-1 expression and decrease epidermal pigmentation (Kim et al. 2019). In summary, while the impact of cellular aging on skin aging remains a topic of ongoing investigation, there are shreds of evidence to suggest that the accumulation of senescent cells could contribute to age-related skin alterations. Therefore, targeting cellular senescence may be a promising therapeutic strategy for delaying skin aging. Transcriptome analysis in dermatology allows researchers to simultaneously examine the expression patterns of thousands of genes and compare them between young and aged skin samples. This approach helps identify key genes and pathways associated with skin aging. By analyzing changes in gene expression, researchers can gain insights into the biological processes that contribute to skin aging, such as alterations in collagen production (Austin et al. 2021), antioxidant defenses (Liu et al. 2021), inflammatory responses (Shehwana et al. 2021), and cellular senescence (Casella et al. 2019). Research that screens for new targets for interventions to prevent or reverse skin aging or identifies genes that become dysregulated during aging is an important tool for developing strategies to modulate the expression or activity of those genes, potentially restoring a more youthful gene expression profiles and improving skin health.

Skin development and maintenance are regulated by keratins, collagens, elastin, and filaggrin-associated genes that orchestrate various cellular processes, including cell proliferation, differentiation, apoptosis, extracellular matrix production, and immune responses (Pfisterer et al. 2021). Among the many genetic molecules essential for skin health, the *HOX* gene is also known to play a role in skin development, patterning, and differentiation, helping to determine body axis and direct the formation of various skin appendages such as hair follicles (Awgulewitsch 2003; Wu et al. 2010). *HOX* genes are a family of related genes that play an important role in animal development as a transcription factor with homeobox, known as the DNA site that encodes homeodomain in mammals (Holland 2013). *HOX* genes regulate body patterns by specifying the identity of embryonic cells and active during embryonic development and play a role in determining the location and specific characteristics of body parts (Pearson et al. 2005). Mutations in these genes can lead to various developmental disorders in the skeletal system (Quinonez et al. 2014). *HOX* genes that exist in humans are divided into four families: A, B, C, and D. A total of 39 *HOX* genes are present in humans, with 11 *HOXA* families on chromosome 7, 10 *HOXB* families on chromosome 17, nine *HOXC* families on chromosome 12, and nine *HOXD* families on chromosome 2. Also, *HOX* genes are divided into anterior-posterior *HOX* genes based on the navel (Morgan 2006). Anterior *HOX* genes are involved in the development of areas close to the navel, such as the head and chest, during early embryogenesis. Despite the fact that the Hox gene family is an essential regulator of organ development and maintenance, the role of *HOX* genes in skin aging and the underlying molecular mechanisms for antiaging interventions remain poorly understood. Thus, Further research is needed to fully elucidate the specific mechanisms through which *HOX* genes influence skin aging and to explore their potential as targets for anti-aging therapies.

In this study, we conducted RNA-Seq analysis to assess the differentially expressed genes (DEGs) associated with skin aging. We specifically examined the transcriptional profiles of human epidermal keratinocyte (HEK) and human dermal fibroblast (HDF) cells derived from neonatal and old age individuals. By comparing the HEK and HDF cells, we observed cell type-specific changes as well as aging-related alterations in both cell types. To identify biologically significant genes, we performed functional analysis and annotated the underlying mechanisms involved in aging. Notably, our focus centered on the expression patterns of *HOX* genes, which have been implicated in skin development and maintenance. Particularly, we detected dynamic transcriptional changes in the regulation of anterior *HOX* genes within the older dermal fibroblast group. Through our investigation, we aimed to explore the correlation between anterior *HOX* genes and skin aging, providing insights into their potential as novel markers of skin aging. The findings of this study shed light on the intricate relationship between *HOX* genes and the aging process in the skin, stimulating a discussion regarding their utility as indicators of skin aging.

## RESULTS

### Transcriptome sequencing analysis results

To explore the transcriptional differences between neonatal and old groups (50s) in human epidermal keratinocytes (HEKs) and human dermal fibroblasts (HDFs), a total of 60 skin samples (every 30 samples per group) were subjected to RNA-Seq analysis. Using an Illumina NovaSeq, an average of 36.5 million raw reads were produced, with a read length of 101 bp. The raw data was qualified through the quality control step for sequence data, and the average number of 36.5 million reads, according for 94.9 % of raw data, were uniquely mapped on the Human Reference Genome (GRCh38) (**Supplementary Table S1**). After gene annotation using the RSEM v1.3.3, among 60,649 genes, a total of 22,621 were commonly expressed in the entire group. We confirmed the uniform data conditions in the gene expression distribution for each sample (**Supplementary Table S2**). To emphasize the associations between samples of each group, the reproducibility of technical replication using PCA plot, density plot, box plot, and pairwise correlation analysis was confirmed based on the overall gene expression in the sample. As a result, similar distributions within the group appeared in solid conditions, and distinct differences between groups were confirmed (**Supplementary Figure S1**). To identify the differentially expressed genes (DEGs) in two skin cell types (HEKs and HDFs) under different aging conditions (neonatal and old), gene expression data from each group were analyzed using DESeq2 (v 1.38.1). The all significant DEGs for each comparison were selected based on a strict cut-off criteria: log2FC >= 1 or log2FC <= –1 (absolute log2FC greater than or equal to 1) and a p-value < 0.05.

### Global landscape of genes expression differences in the HEK and the HDFs

First, for identifying global expression differences between the neonatal and old groups, we explored DEGs and their functions through Metascape analysis using Gene Ontology (GO) and Reactome database. In the comparative analysis of the neonatal and old groups, we identified 1373 DEGs (604 upregulated in the old group and 769 in the neonatal). In these two dramatically aged cell types, we were able to find that they exhibited distinct transcriptomic mosaicism patterns and reviewed their respective function. Most of 604 upregulated DEGs in the old group were highly enriched in the skeletal system morphogenesis (GO:0048705) and signaling receptor regulator activity (GO:0030545). The top 16 function-related genes were listed as *BMP2*, *BMPR1B*, *COMP*, *CXCL8*, *EGR2*, *GDNF*, *GJA1*, *HAS2*, *HTR2B*, *ITGA8*, *ITGB3*, *LIF*, *PKD2*, *RET*, *STC1* and *TBX1* was most highly associated with skeletal system morphogenesis in old group. In addition, the top 10 genes, such as *BMP2*, *CCL20*, *CNTF*, *CXCL8*, *GDNF*, *IL6*, *ITGB3*, *LIF*, *SCG2* and *STC1* was most highly associated with signaling receptor regulator activity in old group. To explore the biological pathway and reactions occurring more specifically in the neonatal and old groups, we analyzed genes with expression differences specific to each group, using the Reactome pathway. In old group, 59 out of 604 were mainly related to the activation of anterior *HOX* genes in hindbrain development during early embryogenesis (R-HSA-5617472). The top 11 function-related genes were listed as *CAMK2A*, *CXCL8*, *H3C6*, *H3C10*, *H3C14*, *IL6*, *MMP3*, *PRKG2*, *RIPK3*, *SAA1* and *TTR* (**Figure 1A and Supplementary Table S3).**

**Figure 1.**
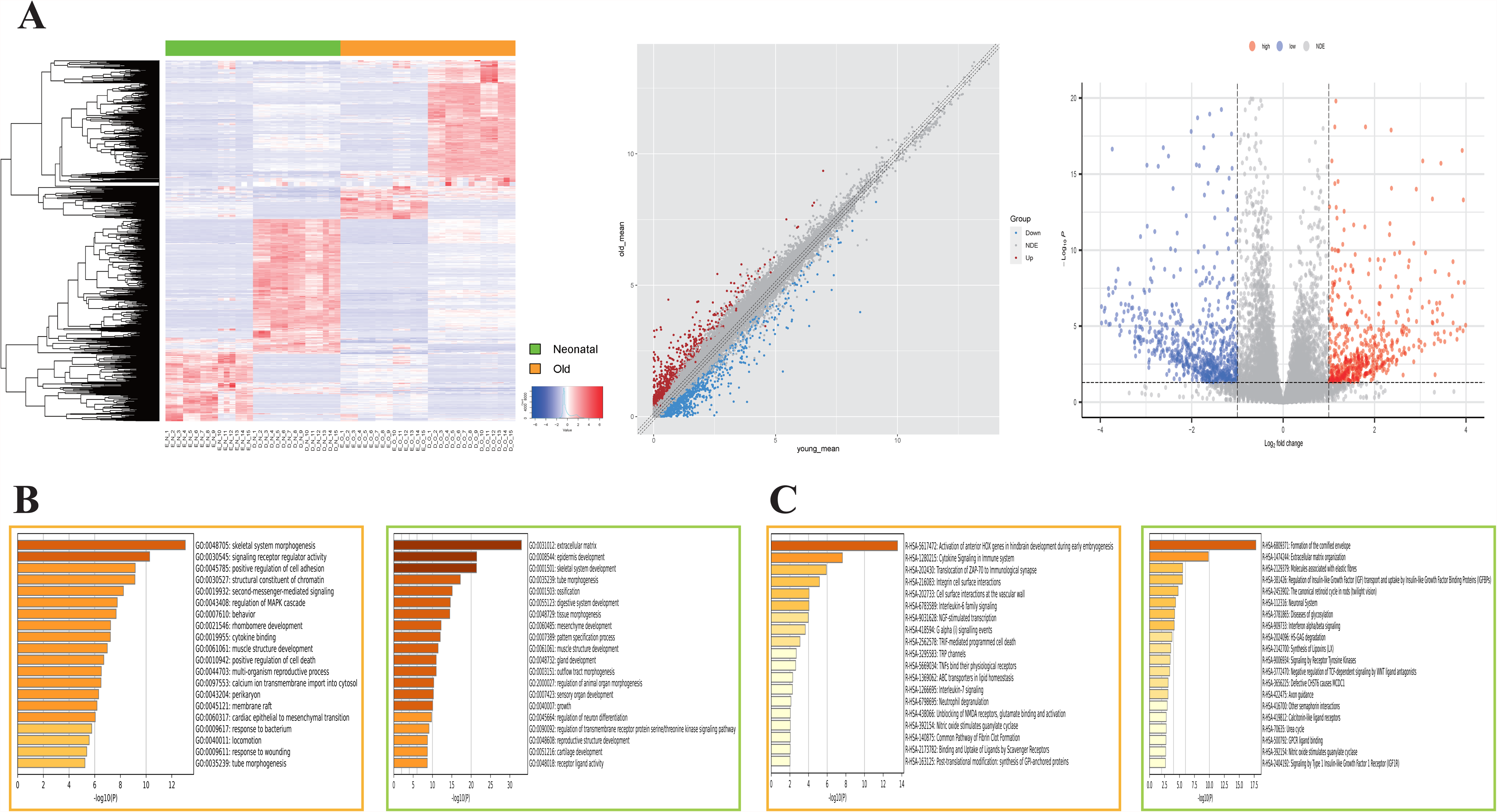
Function prediction analysis for globally DEGs in neonatal and old using Metascape and Reactome pathway. (A) 1373 DEGs (604 upregulated in the old group and 769 upregulated in the neonatal group) were visualized using hierarchical heatmap, volcano and scatter plot. In heatmap, there is a histogram in the color key showing the number of values within each color bar. In volcano and scatter plot, red points with log_2_ fold-change (FC) more than 1 and P-value less than 0.05 were considered statistically upregulated in the old group. Conversely, blue points DEGs with log_2_ fold-change (FC) less than –1 and P-value less than 0.05 were considered downregulated in the old group. Gray points were considered not differentially expressed. (B) There are two color boxes. Orange box shows GO result of upregulated 604 DEGs in the old groups. Green box shows GO result of upregulated 769 DEGs in the neonatal groups. (C) Orange box shows Reactome pathway result of upregulated 604 DEGs in the old groups. Green box shows Reactome pathway result of upregulated 769 DEGs in the neonatal groups.

While in neonatal group, 769 upregulated DEGs were highly enriched in the extracellular matrix (GO:0031012), epidermis development (GO:0008544) and skeletal system development (GO:0001501). 86 out of 769 were mainly related to the extracellular matrix. Fourteen of the 86 genes, including *COL1A1*, *COL5A1*, *COL11A1*, *FGF10*, *FOXC1*, *FOXC2*, *FOXF1*, *FOXF2*, *GPC3*, *NTN1*, *PRICKLE1*, *SFRP2*, *WNT2* and *WNT5A,* were involved in extracellular matrix function as well as various functions involved in multicellular organism development. Nineteen genes including *BCL2*, *BMP4*, *FGF10*, *FGF9*, *FOXC1*, *FOXC2*, *FOXF1*, *GATA6*, *GLI2*, *HOXA13*, *ID2*, *PRICKLE1*, *SFRP2*, *SIX1*, *SIX2*, *SOX11*, *TCF21*, *WNT2*, and *WNT5A*, known to be key players in mesenchymal development were enriched in the skeletal system development and the epidermis development. In Reactome pathway analysis, 28 out of 769 were mainly related to the formation of the cornified envelope (R-HSA-6809371). The 28 genes listed as *8* keratins (*KRT*), 9 late cornified envelope (*LCE*), 4 Small proline-rich proteins (SPRR) gene family, *CDSN*, *KLK12*, *KLK13*, *LIPK*, *LORICRIN*, *PCSK6*, and *TGM1* (**Figure 1 and Supplementary Table S3)**.

### Independent Transcriptomic Changes in the HEKs and HDFs with Skin Aging

Metascape GO functional prediction for gene expression changes with age in HEKs and HDFs, we have observed a series of significant biological alterations, as illustrated in **Figure 2**. In the case of HEKs, we have identified 917 DEGs (311 in the old HEKs and 606 upregulated genes in the neonatal HEKs). The functional categorization of these DEGs has provided valuable insights. Specifically, we found that 27 out of the 311 upregulated genes in the old HEKs were primarily associated with the ameboidal-type cell migration (GO:0001667). Furthermore, 46 out of the 311 upregulated genes were prominently associated with skin development (GO:0043588) in the old HEKs. Top 10 genes (*ITGA4*, *CYP1B1*, *TGFBR3*, *FN1*, *CLDN1*, *JAM3*, *WNT5B*, *EREG*, *COL5A1*, and *BMPR1B*) are shown close relations with GO functions in the old HEKs. We also examined the Reactome pathway (https://reactome.org) to gain further insights. The analysis of DEGs revealed that in the old HEKs, 17 out of the 311 upregulated genes were primarily associated with Extracellular matrix organization (R-HSA-1474244) and Formation of the cornified envelope (R-HSA-6809371). Among these, the higher associated genes with pathways were *ADAMTS1*, *COL5A1*, *COL5A2*, *COL5A3*, *FN1*, *SPP1*, and *THBS2*, demonstrating a strong contribution in skin homeostasis and remodeling in the old HEKs. Additionally, 10 genes (*CASP14*, *DSG1*, *KRT2*, *KRT31*, *KRT6B*, *KRT6C*, *KRT75*, *KRT79*, *LCE3C*, and *TCHH*) were predominantly linked to the Formation of the cornified envelope pathway (R-HSA-6809371) in the old HEKs. As for neonatal HEKs, we observed that 80 and 40 out of the 606 downregulated genes were primarily linked to the epidermis development (GO:0008544) and the external encapsulating structure (GO:0045229). The top 10 genes highly associated with those function was *TREX1*, *SMPD3*, *CCN2*, *HMGB2*, *LGALS9*, *ESR1*, *CTSH*, *ARG1*, *NR4A1*, and *PARP9*.

**Figure 2.**
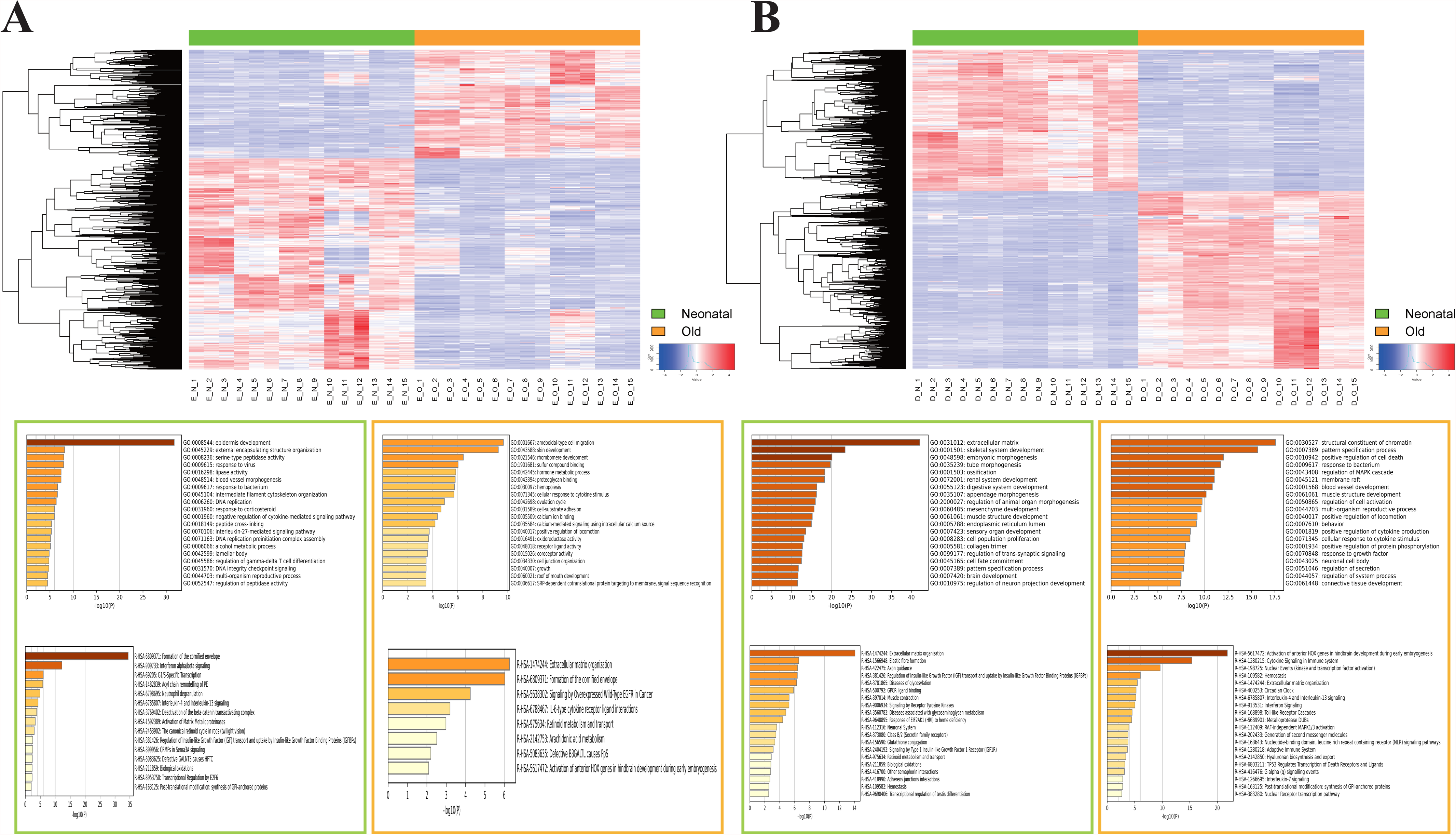
Age-dependent DEG and their functions in skin cell types (HEKs and HDFs). (A) Hierarchical clustering heatmap for 917 DEGs (311 upregulated in the old the HEK and 606 upregulated in the neonatal the HEK) are represented. There is a histogram in the color key showing the number of expression values within each color bar. (B) Hierarchical clustering heatmap for 1953 DEGs (1092 upregulated in the old the HDFs and 861 upregulated in the neonatal the HDFs) are represented. The green boxes below the heatmap are the GO and Reactome pathway results of upregulated genes in the neonatal groups. The orange boxes below the heatmap are the GO and Reactome pathway results of upregulated genes in the old groups.

Through Reactome pathway analysis, we observed that 39 genes, including 10 keratins, 11 late cornified envelope protein, 9 Small proline-rich proteins (*SPRR*) gene family, exhibited the highest association with the formation of the cornified envelope pathway (R-HSA-6809371) in the neonatal HEKs. These findings provide valuable insights into the molecular differences occurring with age in HEKs, highlighting the significant roles of the ameboidal-type cell migration, skin development, and epidermis development in the aging process (**Figure 2A and Supplementary Table S4**). Both HEKs from the old and neonatal groups showed expression of key genes involved in skin development and extracellular matrix organization, and here we examined the independent transcriptome expression in each aging group and found that the neonatal HEKs showed more transcriptomic activity related to epidermis development.

In the case of HDFs, we identified 1953 DEGs (1092 in old HDF and 861 upregulated genes in the neonatal HDFs). Functional categorization of DEGs revealed interesting insights into the gene expression profiles of the old HDFs. Among the 1092 DEGs, 69 were primarily associated with the structure constituent of chromatin (GO:0030527). The top 10 genes, namely *ADRA2A*, *ATF3*, *BCL6*, *EP300*, *EPAS1*, *HOXA5*, *HOXA7*, *INPP5D*, *NR4A1*, and *TLR4*, showed the highest association with the structure constituent of chromatin in the old HDFs. Furthermore, 136 upregulated DEGs, including early growth response (EGR) protein in old HDFs were related to the pattern specification process (GO:0007389). Regarding Reactome pathway analysis, 171 out of the 1092 DEGs were prominently associated with the “Activation of anterior HOX genes in hindbrain development during early embryogenesis” pathway (R-HSA-5617472). The top 12 genes highly linked to this pathway in the old HDFs were *BCL2L1*, *CAMK2A*, *CTSK*, *CTSV*, *FOS*, *ITPR3*, *MMP3*, *PLCG2*, *PRKCD*, *RORA*, *RPS6KA1*, and *STAT1*. Neonatal HDFs exhibited three distinct functional signatures, as shown in the gene ontology (GO) results. Among 861 DEGs, 101 were primarily associated with the extracellular matrix (GO:0031012). The function-related genes with the extracellular matrix in the neonatal HDFs included *COL11A1*, *COL3A1*, *CTHRC1*, *FGF10*, *FOXC1*, *FOXC2*, *FOXF1*, *GPC3*, *PRICKLE1*, *SFRP2*, *SOX9*, *WNT2*, and *WNT5A* which are a vital gene network in tissue morphogenesis and development (Yu et al. 2021). Sixty-four and 96 out of 861 DEGs were predominantly related to skeletal system development (GO:0001501) and embryonic morphogenesis (GO:0048598), respectively. Collagen, fibroblast growth factor (*FGF*), forkhead Box C (*FOX*), SIX Homeobox (*SIX*), homeobox (*HOX*), and Sry-type HMG box (*SOX*) genes were enriched at both biological function categories. In additional examination with Reactome, 44 and 10 out of 861 DEGs in the neonatal HDFs were primarily associated with the process of extracellular matrix organization (R-HSA-1474244) and Elastic fibre formation (R-HSA-1566948), respectively. (**Figure 2B and Supplementary Table S5**) Overall, the findings suggest that aging in HEKs is associated with ameboidal-type cell migration, skin development, and extracellular matrix organization, while in HDFs, aging is linked to changes in chromatin structure, pattern specification, and activation of *HOX* genes. Additionally, neonatal HDFs showed distinct functional signatures related to the extracellular matrix, skeletal system development, and embryonic morphogenesis.

### Activation of 39 *HOX* genes in HEKs and HDFs

The combined analysis of HEK and HDF cells in terms of specificity and age-specific transcriptomic expression differences revealed valuable insights into the molecular biological processes occurring in skin cells. One notable finding was a drastic change observed in the *HOX* gene family expression in HDFs, with opposite gene expression patterns between the neonatal and old groups. Thus, we observed a total of 39 *HOX* genes which are known as a group of genes that encode transcription factors involved in embryonic development and pattern formation and organized into four clusters: *HOXA*, *HOXB*, *HOXC*, and *HOXD*. Interestingly, the HDF cells type, on the other hand, showed a dramatic 25 upregulated and 10 downregulated changes relative to old HDFs. In addition, *HOXC4* and *HOXC10* genes were expressed in both the neonatal and old HDFs but have a little higher gene expression in the old HDFs (**Figure 3A**). To make it easier to see the distinctly different expression patterns of 39 *HOX* genes in the HDFs, a schematic gene expression alteration in the neonatal and old HDFs was drawn. Almost all anterior *HOX* genes which involved in the development of structures and tissues in the head and anterior regions of the body, including the brain, face, sensory organs, and cranial nerves, were upregulated in the old HDFs. In contrast, most of posterior *HOX* genes which has crucial role in the development of structures and tissues in the posterior part of the body, including the spinal cord, limbs, urogenital system, and tail, were downregulated in the old HDFs (Mallo et al. 2010). In the neonatal HDFs, the opposite of the expression patterns was shown (**Figure 3B**). Only four of the 39 *HOX* genes (*HOXA1*, *HOXA9*, *HOXC12*, and *HOXC13)* showed HEK cell type-specific expression when compared to HDFs, while there was no difference in *HOX* gene expression between the neonatal and old groups. Seven of the 35 differentially expressed *HOX* genes in the old HDFs (*HOXA2*, *HOXA3*, *HOXA4*, *HOXA5*, *HOXA6, HOXA7*, and *HOXC11*) were also expressed in the neonatal and old HEKs. The *HOXA2*, *HOXA3*, *HOXA4*, *HOXA5*, *HOXA6* and *HOXA7* genes were shown higher expressions in the neonatal HEKs (**Figure 3A)**.

**Figure 3.**
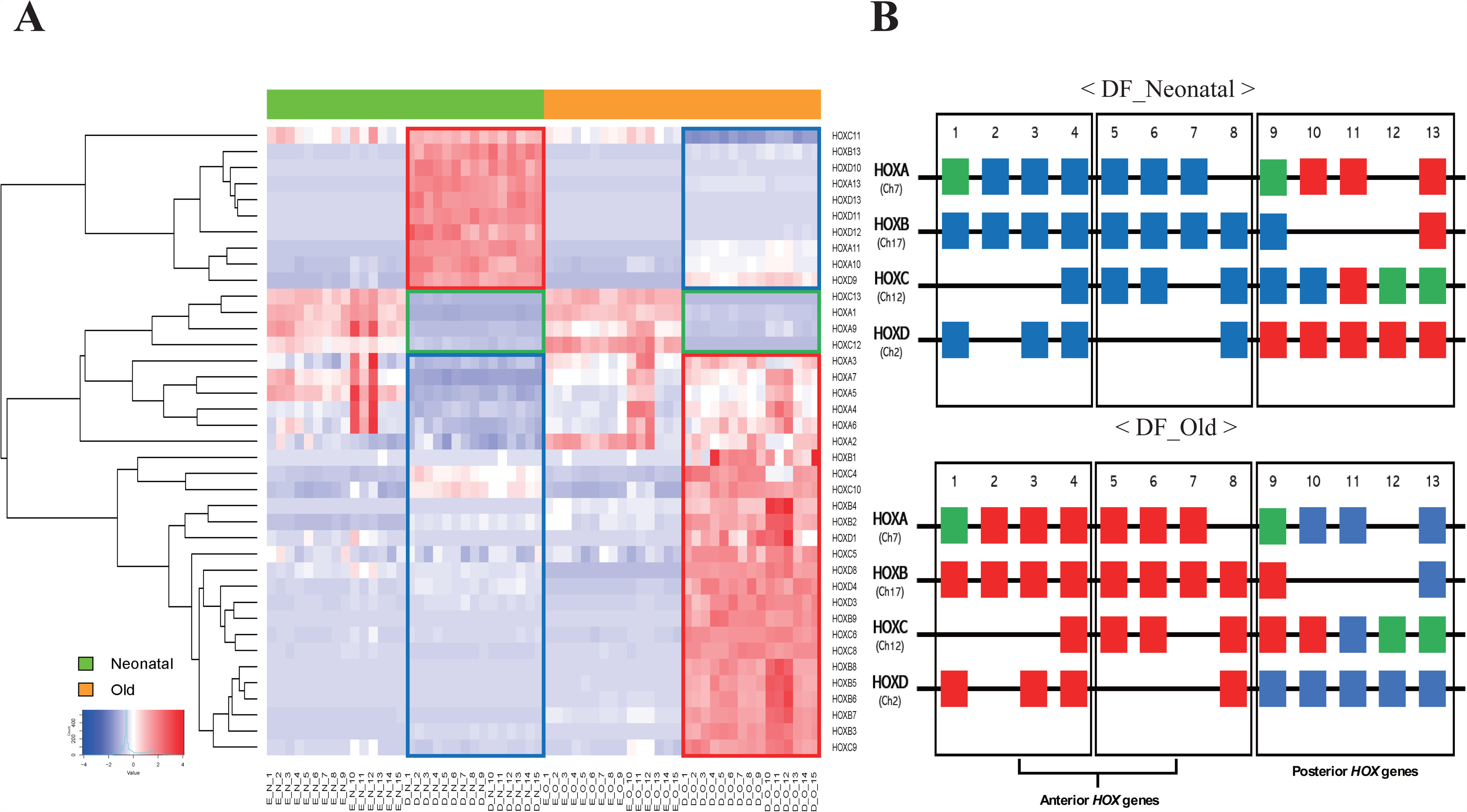
Expression of 39 *HOX* genes in the HEK and the HDFs between neonatal and old groups. (A) 39 *HOX* genes were visualized using heatmap. There is a histogram color key showing the number of value within each color bar. Red boxes are clustering of upregulated *HOX* genes in the the HDFs of each group. Blue boxes are clustering of downregulated *HOX* genes in the the HDFs of each group. Green boxes are clustering of commonly downregulated *HOX* genes in the the HDFs both groups. (B) We drawn two schematics that easier to see different expression patterns of *HOX* genes in the the HDFs. Red boxes mean the *HOX* genes that were upregulated in each group. Blue boxes mean the *HOX* genes that were downregulated in each group. Green boxes mean the *HOX* genes that were commonly not expressed in both groups.

In addition to the previously mentioned findings, the analysis revealed additional insights into the gene expression patterns and regulatory factors associated with age-related changes in HDFs. Specifically, transcriptional cofactor genes, *EGR2* and *RARB* were found to be highly expressed in old HDFs. These genes are known to play a role in the transcriptional activation of *HOX* genes (Serpente et al. 2005; Addison et al. 2018). The expression levels of eight histone proteins, which contribute to DNA condensation and play a crucial role in regulating gene expression, were found to be higher in old HDFs compared to other groups (**Figure 4**). These findings collectively support the notion that the activation of various transcription factors, including transcriptional cofactors and histone proteins, is likely to occur in old HDFs. This implies a complex regulatory network underlying the age-related changes in gene expression patterns. Our study firstly identified age-dependent expression differences in anterior-posterior *HOX* genes in HDFs. While HEKs showed similar expression patterns of *HOX* genes regardless of age, HDFs exhibited clear age-dependent expression differences. Specifically, the upregulation of anterior *HOX* genes and downregulation of posterior *HOX* genes were observed with aging. These findings provide further insights into the molecular mechanisms underlying age-related changes in gene expression and highlight the dynamic regulation of *HOX* genes in the aging process.

**Figure 4.**
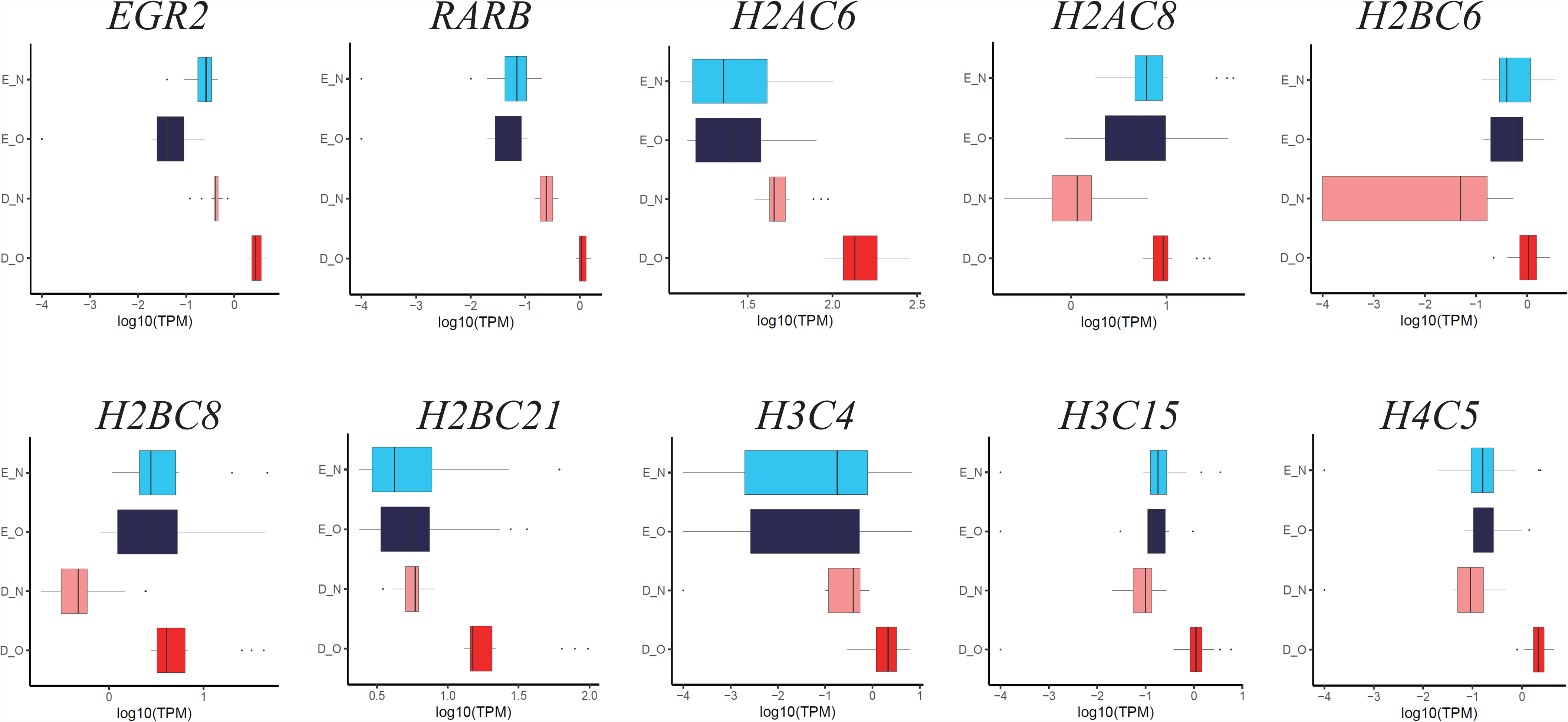
Differentially expression of 13 genes associated activation of anterior *HOX* genes in hindbrain development during early embryogenesis. 13 out of 91 (related to activation of anterior *HOX* genes in hindbrain development during early embryogenesis) genes overlapped our data. Bar graphs show the values of 13 genes age-dependent expression in the the HEKs and the HDFs. Light blue bars indicate expression values within the neonatal the HEK. Navy blue bars indicate expression values within the old the HEK. Pink bars indicate expression values within the neonatal the HDFs. Red bars indicate expression values within the old the HDFs. All expression values (TPM) were normalized to log_10_ (TPM).

### Age-dependent Differentially Expression of Skin Barrier-related Factors

The decrease of collagen and elastin fibers is a typical response and bio indicator of skin aging (Taszkun et al. 2019; Baumann et al. 2021). Thus, we also investigated the representative component of the skin that plays a significant role in maintaining its health and integrity: collagen type I (*COL1A1*, *COL1A2*), collagen type III (*COL3A1*), collagen type lJ (*COL5A1*, *COL5A2*, *COL5A3*), and elastin fibers (*ELN*, *FBLN1*, *FBLN2*) (**Figure 5A**). As shown in Figure 5A, all nine genes had age-dependent different expression patterns in the HEKs and the HDFs and had higher gene expression activities in the HDFs compared to the HEKs. In HEK cells, *COL1A1* showed similar expression values between neonatal and old groups. However, in HDF cells, *COL1A1* had higher expression values in neonatal groups compared to old groups. For *COL1A2*, HEK cells exhibited higher expression values in the old groups, while HDF cells showed higher expression values in the neonatal groups. *COL3A1* had very low expression values in both neonatal and old groups in HEK cells, but slightly higher expression values in the old groups. In HDF cells, *COL3A1* had higher expression values in neonatal groups compared to old groups. In HEK cells, both *COL5A1* and *COL5A2* had higher expression values in the old groups, whereas in HDF cells, these genes had higher expression values in the neonatal groups. Regarding *COL5A3*, it showed higher expression values in the old groups in HEK cells, while in HDF cells, there were similar expression values between neonatal and old groups. In HEK cells, *ELN* did not show significant expression values, whereas in HDF cells, it exhibited notable expression. Additionally, the neonatal groups had higher expression values of *ELN* compared to the old groups in HDFs. Both *FBLN1* and *FBLN2* exhibited similar expression patterns in both HEK and HDF cells. The neonatal groups had higher expression values than the old groups for both genes, and this difference was more pronounced in HEK cells.

**Figure 5.**
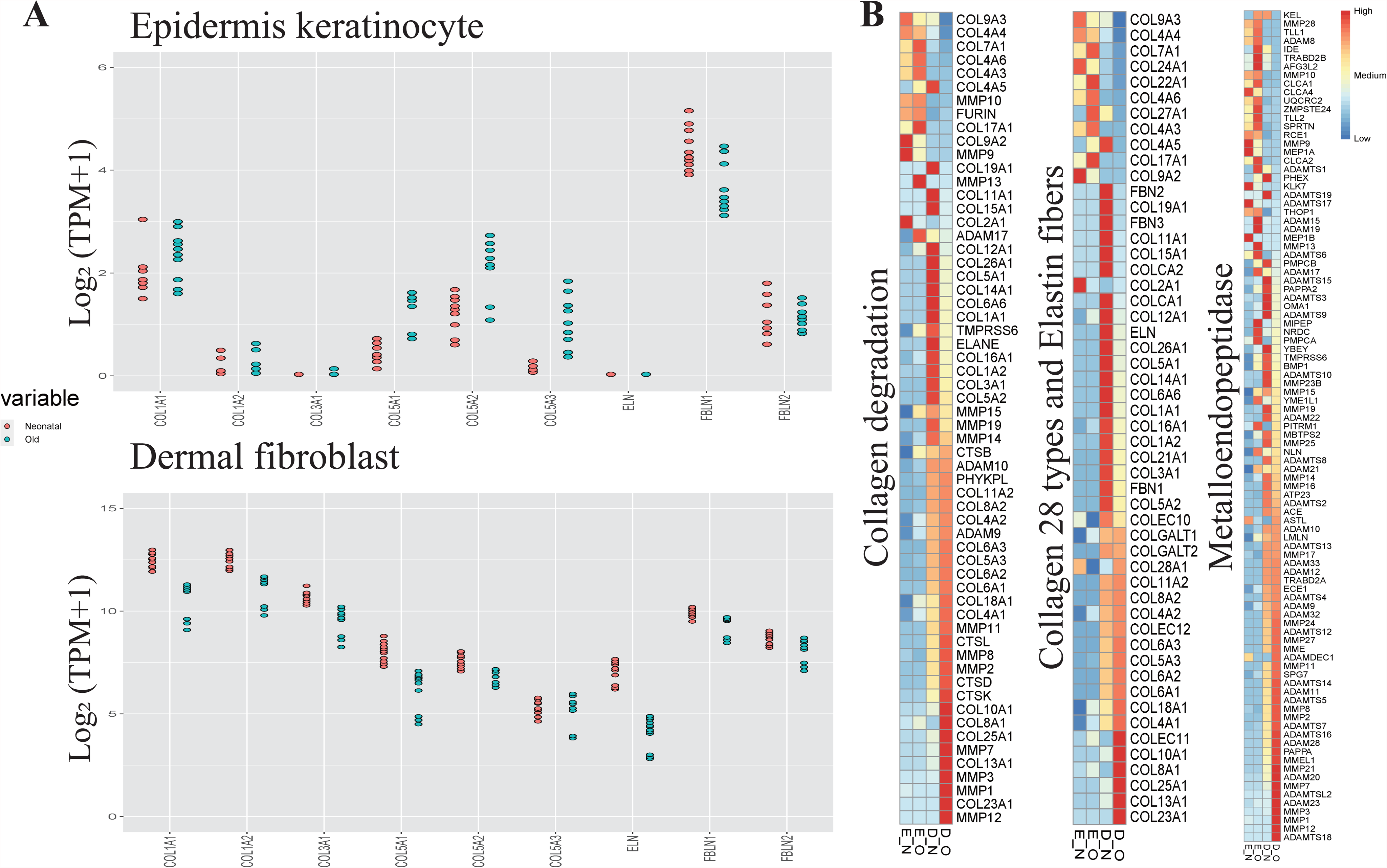
Differentially expression of various skin barrier-related factors including collagen and elastin fibers. (A) Dot plots show the nine genes (*COL1A1*, *COL1A2*, *COL3A1*, *COL5A1*, *COL5A2*, *COL5A3*, *ELN*, *FBLN1*, *FBLN2*) expression in paired neonatal and old in the the HEK and the HDFs. Red dots indicates expression values of the neonatal groups. Green dots indicates expression values of the old groups. All expression values (TPM) were normalized to log_2_ (TPM+1). (B) Heatmap plots of differentially expressed genes from the three categories. Three heatmap plots show the expression patterns of the genes in each category. In the collagen degradation, 60 genes show patterns of age-dependent differentially expression in the the HEK and the HDFs. In the collagen 28 types and elastin fibers, 52 genes show patterns of age-dependent differentially expression in the the HEK and the HDFs. In the metalloendoipeptidase, 97 genes show patterns of age-dependent differentially expression in the the HEK and dermal fibrobalsts. There is color key showing the number of expression values besides heatmap plots. High expression is indicated in red and low expression is indicated in blue. All expression values (TPM) were normalized to Z score.

We explored the expression levels of all genes involved in collagen synthesis and collagen degradation, including collagen degradation, collagen 28 types and elastin fibers, and metalloendopeptidase, in the two cell types by age. (**Figure 5B**). Our analysis revealed significant age-dependent expression differences in 60 genes related to collagen degradation in both HEK and HDF cells. In HEK cells, 10 of these genes, including *COL4A3*, *COL4A4*, *COL4A6*, *COL7A1*, *COL9A2*, *COL9A3*, *COL17A1*, *FURIN*, *MMP9*, and *MMP10*, showed higher upregulation. In neonatal HDFs, 4 genes (*COL4A5*, *COL11A1*, *COL15A1*, and *COL19A1*) exhibited high expression levels. *ADAM17* and *MMP13* were highly expressed in old HEK cells, while *COL2A1* was highly expressed in neonatal HEK cells. In the HDFs, 43 out of the 60 genes demonstrated clear age-dependent expression differences. Among these, 15 genes (*COL1A1*, *COL1A2*, *COL3A1*, *COL5A1*, *COL5A2*, *COL6A6*, *COL12A1*, *COL14A1*, *COL16A1*, *COL26A1*, *ELANE*, *MMP14*, *MMP15*, *MMP19*, and *TMPRSS6*) were highly expressed in neonatal HDFs. On the other hand, 28 genes (*ADAM9*, *ADAM10*, *COL4A1*, *COL4A2*, *COL5A3*, *COL6A1*, *COL6A2*, *COL6A3*, *COL8A1*, *COL8A2*, *COL10A1*, *COL11A2*, *COL13A1*, *COL18A1*, *COL23A1*, *COL25A1*, *CTSB*, *CTSD*, *CTSK*, *CTSL*, *MMP1*, *MMP2*, *MMP3*, *MMP7*, *MMP8*, *MMP11*, *MMP12*, and *PHYKPL*) showed higher expression in the old HDFs.

For the analysis focused on genes related to collagen (28 types) and elastin fibers, 9 out of 52 genes (*COL4A3*, *COL4A4*, *COL4A6*, *COL7A1*, *COL9A2*, *COL9A3*, *COL17A1*, *COL22A1*, and *COL24A1*) showed higher upregulation in the HEKs. *COL4A5* and *COL27A1* exhibited higher expression in the old HEK cells and neonatal HDFs. In the HDFs, 41 genes demonstrated age-dependent expression differences. Among these, 21 genes (*COL1A1*, *COL1A2*, *COL3A1*, *COL5A1*, *COL5A2*, *COL6A6*, *COL11A1*, *COL12A1*, *COL14A1*, *COL15A1*, *COL16A1*, *COL19A1*, *COL21A1*, *COL26A1*, *COLCA1*, *COLCA2*, *COLEC10*, *ELN*, *FBN1*, *FBN2*, and *FBN3*) were highly expressed in the neonatal HDFs. Conversely, 18 genes (*COL4A1*, *COL4A2*, *COL5A2*, *COL6A1*, *COL6A2*, *COL6A3*, *COL8A1*, *COL8A2*, *COL10A1*, *COL11A2*, *COL13A1*, *COL18A1*, *COL23A1*, *COL25A1*, *COLEC11*, *COLEC12*, *COLGALT1*, and *COLGALT2*) showed higher expression in old HDFs. *COL2A1* was most highly expressed in neonatal HEK cells, while *COL28A1* exhibited similar expression values in the neonatal HEK cells and old HDFs.

In 97 out of 111 genes related to metalloendopeptidase activity, 7 genes (*ADAMTS17*, *CLCA4*, *KLK7*, *MEP1A*, *MEP1B*, *MMP9*, and *RCE1*) were more upregulated in the neonatal HEK cells. In the old HEKs, 27 genes (*ADAM8*, *ADAM15*, *ADAM17*, *ADAM19*, *ADAM21*, *ADAMTS1*, *ADAMTS6*, *AFG3L2*, *CLCA1*, *CLCA2*, *IDE*, *MIPEP*, *MMP10*, *MMP13*, *MMP28*, *NLN*, *NRDC*, *PITRM1*, *PMPCA*, *SPRTN*, *THOP1*, *TLL1*, *TLL2*, *TRABD2B*, *UQCRC2*, *YME1L1*, and *ZMPSTE24*) showed higher upregulation. In the neonatal HDFs, 25 genes (*ACE*, *ADAM22*, *ADAMTS2*, *ADAMTS3*, *ADAMTS8*, *ADAMTS9*, *ADAMTS10*, *ADAMTS15*, *ADAMTS19*, *ATP23*, *BMP1*, *KEL*, *MBTPS2*, *MMP14*, *MMP15*, *MMP16*, *MMP19*, *MMP23B*, *MMP25*, *OMA1*, *PAPPA2*, *PHEX*, *PMPCB*, *TMPRSS6*, and *YBEY*) exhibited higher expression levels. In the old HDFs, 38 genes (*ADAM9*, *ADAM10*, *ADAM11*, *ADAM12*, *ADAM20*, *ADAM23*, *ADAM28*, *ADAM32*, *ADAM33*, *ADAMDEC1*, *ADAMTS4*, *ADAMTS5*, *ADAMTS7*, *ADAMTS12*, *ADAMTS13*, *ADAMTS14*, *ADAMTS16*, *ADAMTS18*, *ADAMTSL2*, *ASTL*, *ECE1*, *LMLN*, *MME*, *MMEL1*, *MMP1*, *MMP2*, *MMP3*, *MMP7*, *MMP11*, *MMP12*, *MMP17*, *MMP21*, *MMP24*, *MMP27*, *PAPPA2*, *SPG7*, and *TRABD2A*) showed higher upregulation. These findings not only provide valuable insights into the expression patterns of collagen and metalloendopeptidase-related genes in different cell types and age groups, but also provide a blueprint for cell-specific molecular markers needed in many molecular biology studies for skin improvement and maintenance.

## DISCUSSION

The role of *HOX* genes in embryonic development and their influence on cell fate determination along the anterior-posterior body axis have been well-established (Rux and Wellik 2017). However, their involvement in skin aging has not been extensively studied. To strengthen the reliability of our results, we explored previous research on the functions of *HOX* genes in skin wound healing, skeletal development, and individual gene functions. Danielle R. Rux demonstrated that the activation of the *HOXA11* gene has a significant impact on the differentiation of mesenchymal stem cells into osteoblasts. This finding led us to hypothesize that down-regulation of *HOXA11*, a posterior *HOX* gene, could potentially impair the differentiation ability of mesenchymal stem cells. Additionally, through our investigation of individual functions of 39 *HOX* genes and other relevant literature, we came across *HOXA7* and *HOXB13* genes.

The expression of *HOXA7* has been found to exhibit an inverse relationship with keratinocyte differentiation (La Celle and Polakowska 2001). When *HOXA7* is downregulated, the activity of keratinocyte differentiation is enhanced, while upregulation of *HOXA7* inhibits keratinocyte differentiation. Keratinocyte differentiation is crucial for the formation of the stratum corneum, which helps maintain skin health by preserving external protection and moisture levels (de Farias Pires et al. 2016). In **Figure 1C**, we observe that the formation of the cornified envelope is the most significant biological pathway in the neonatal HDFs group. Furthermore, **Figure 3B** reveals that *HOXA7* expression is downregulated in the neonatal HDFs group and upregulated in the old HDFs group. *HOXB13* gene expression has been associated with fetal skin development (Kömüves et al. 2003) and the wound healing process in fetal skin (Stelnicki et al. 1998). Conversely, degradation of *HOXB13* reduces the regulation of epidermal differentiation in adult skin (Mack et al. 2003). Interestingly, *HOXA7* belongs to the anterior *HOX* genes, while *HOXA11* and *HOXB13* belong to the posterior *HOX* genes. We observed that *HOXA7* gene expression was downregulated, and *HOXA11* and *HOXB13* gene expression were upregulated in the neonatal HDFs group. Conversely, *HOXA7* gene expression was upregulated, and *HOXA11* and *HOXB13* gene expression were downregulated in the old HDFs group. These findings provide confidence that age-dependent expression differences of anterior-posterior *HOX* genes contribute to skin aging by influencing the differentiation of mesenchymal stem cells.

Skin aging is a complex process characterized by a decline in collagen and elastin fibers, resulting in prominent signs such as wrinkles and reduced elasticity (Tzaphlidou 2004; Haeusler 2015). The HDFs play a crucial role in this aging process as they are responsible for collagen synthesis (Garner and surgery 1998). and are closely associated with skin aging (Lago and Puzzi 2019). The decrease in collagen synthesis is a well-established hallmark of skin aging and contributes to the development of wrinkles and diminished skin elasticity (Wang and Dreesen 2018). The HDFs, which are abundant in the dermal layer, play a vital role in maintaining collagen homeostasis and overall skin health. In our study, we specifically examined collagen type I (*COL1A1*, *COL1A2*), collagen type III (*COL3A1*), and collagen type V (*COL5A1*, *COL5A2*, *COL5A3*), which are prominent collagen types in the skin (Zhang et al. 2023). Additionally, we investigated the expression specificity of elastin fibers, including *ELN*, *FBN1*, and *FBN2*. Collagen and elastin fibers are vital components that contribute to the structural integrity, elasticity, and resilience of the skin (Heinz 2021). Understanding the regulation and expression of these collagen and elastin fibers is critical for comprehending the mechanisms underlying skin health and aging. Further investigation into their dynamic changes across the skin cell types and tissues can provide valuable insights for developing interventions to enhance skin quality and combat the signs of aging.

Our findings revealed significant age-dependent expression differences in various factors related to skin barrier function, as depicted in **Figure 5B**. Among these factors, MMPs exhibited notable expression changes in both the HEKs and the HDFs. Specifically, we identified a cluster of genes that were highly upregulated in the old HDFs within three categories: collagen degradation, collagen 28 types and elastin fibers, and metalloendopeptidase. Within these categories, we identified 23 genes that were consistently upregulated in the old HDFs, including *ADAM9*, *ADAM10*, *COL4A1*, *COL4A2*, *COL5A3*, *COL6A1*, *COL6A3*, *COL8A1*, *COL8A2*, *COL10A1*, *COL13A1*, *COL18A1*, *COL23A1*, *COL25A1*, *MMP1*, *MMP2*, *MMP3*, *MMP7*, *MMP8*, *MMP11*, and *MMP12*. Notably, *MMP2*, a member of the *MMP* gene family, plays a crucial role in the breakdown of collagen and other extracellular matrix components. In skin, *MMP*s are responsible for collagen degradation, and the balance between collagen breakdown and synthesis is essential for maintaining skin integrity. However, during the aging process, collagen resynthesis becomes less efficient compared to the activity of *MMP*s. This imbalance leads to the loss of skin elasticity, the formation of wrinkles, and other typical signs of skin aging. Given this context, we investigated the potential correlation between *MMP*s and *HOX* genes. Previous research has indicated that *HOXD3* regulates the expression of *MMP2*, suggesting a potential connection between these factors. Understanding the interplay between *MMP*s and *HOX* genes could provide valuable insights into the molecular mechanisms underlying skin aging (Hamada et al. 2001). As shown in **Figure 3**, our results confirmed that *HOXD3* is upregulated in the old HDFs and downregulated in the neonatal HDFs. Considering our findings and previous studies, we are increasingly convinced that the differential expression of anterior-posterior *HOX* genes contributes to the aging process of the skin.

Our study highlights the potential of *HOX* genes as new markers for skin aging and proposes that age-dependent expression differences of *HOX* genes contribute to the aging process of the skin. However, it is important to note that establishing a direct link between the age-dependent expression differences of the 39 anterior-posterior *HOX* genes and skin aging is challenging due to the complex relationship between *HOXD3* and *MMP2*. We hope that our study serves as a starting point for further investigations into new skin aging markers based on genes regulated by differential expression of anterior-posterior *HOX* genes.

## MATERIALS AND METHODS

### Cell culture and Treatment

The primary human the HDFs (HDFs) and primary human epidermal keratinocytes (HEKs) were purchased from PromoCell (Heidelberg, Germany). HDFs and HDFs at passages 3-5 were cultured in a Fibroblast Growth Medium 2 with supplementMix and in and Keratinocyte Growth Medium 2 with supplementMix, respectively. To compare gene expressions during aging, HDFs and HEKs from neonatal and 50s age were used.

### RNA Sample Preparation

The the HEKs and the HDFs were separated and homogenized in 500 µL of TRIzol reagent (Invitrogen, Carlsbad, CA, USA) using a microhomogenizer following the manufacturer’s instructions. Total RNA was isolated from frozen using RNeasy Mini Kit (Qiagen, Hilden, Germany).

### RNA Sequencing Library Construction

Prior to constructing RNA sequencing libraries, the quality of all RNA samples was checked using the 28S/18S ratio and RNA integrity number (RIN) value using an Agilent Bioanalyzer 2100 system (Agilent Technologies, Santa Clara, CA, USA). All RNA samples showed RIN values higher than 9.0. mRNA molecules were enriched and purified from 500 ng of the qualified RNA samples using oligo-dT magnetic beads. Double-stranded cDNA was immediately synthesized by SuperScript III reverse transcriptase (Thermo Fisher Scientific Inc., Waltham, MA, USA). According to the instructions of the TruSeq RNA Sample Prep Kit (Illumina, San Diego, CA, USA), a sequential process of end repair, poly-A addition, and adaptor ligation on both ends was carried out. The processed cDNA libraries were subjected to library enrichment by polymerase chain reaction (PCR) and size selection to the exact appropriate size of fragments using the BluePippin Size-Selection system (Sage Science, Beverly, MA, USA). The final selected libraries were evaluated with an Agilent Bioanalyzer 2100 system and were 400–500 bp in size. The cDNA libraries were sequenced with an Illumina HiSeq2500 (Illumina San Diego, CA, USA), which generated paired end reads of approximately 100 bp in size.

### Data Analysis

#### 1) Quality Control

Raw sequencing data was evaluated to discard low-quality reads by FASTQC (https://www.bioinformatics.babraham.ac.uk) as follows steps: Reads including more than 10% of skipped bases (marked as ‘N’); sequencing reads including more than 40% of bases whose quality score is less than 27; their average quality score (<27). Quality distributions of nucleotides, GC contents, PCR duplication properties, and k-mer sequencing data frequencies were calculated.

#### 2) Read Mapping and Differentially Expressed Genes (DEG) Analysis

High-quality reads mapped on the Human reference genome (Genome sequence, primary assembly(GRCh38)) using aligner STAR v2.7.8a. We only used uniquely mapped read pairs for the analysis of differentially expressed genes (DEGs). To identify DEGs, gene expression count data were generated using RSEM v1.3.3, and the R package for comparing TPM (Transcripts per million) count with the normalization method was used. Differentially expressed genes for 2 groups were analyzed using DESeq2 v1.34.0 methods in R v4.2.2. The DEGs with log2 fold-change (FC) more than 1 and P-value less than 0.05 was considered statistically significant.

#### 3) Data Statistical Analysis and Visualization

For the expression data across all samples, the log_2_ transformed TPM values were represented by qualitative characteristics of normalized data, including the count distributions and variability between biological replicates. The general analysis for statistical validation, including pairwise correlation analysis and scatterplot, the hierarchical clustering heatmaps and volcano plots, and principal components analysis (PCA) plots were performed with the ggplot2 package using R v4.2.2. The heatmap clustering analysis of DEGs was performed based on the log_2_ TPM values, and the heatmap was generated using gplots package v 3.1.3 with the popular clustering distance (euclidean) and hierarchical clustering method (complete) functions.

#### 4) Integrative Function Classification analysis for DEGs

Gene ontology (GO) was performed using Metascape (http://metascape.org/gp/index.html). The Metascape analysis workflow followed these criteria: First, the multiple gene lists identified from DEG analysis were used as input genes. Second, the main categories of gene functions were extracted for the Reactome pathway. Third, functional enrichment analysis was performed with default parameters (min overlap of 3, enrichment factor of 1.5, and P-value of 0.01) for filtering. The Reactome pathway database that interprets biological pathways is also used to identify the functional role of genes that show differences in gene expression depending on aging.

## ACKNOWLEDGEMENTS

The authors gratefully acknowledge Center for Bio-Medical Engineering Core Facility at Dankook University for providing critical reagents and equipment. This project was supported by project COSMAX and COSMAX BTI center of Korea Cosmetic industry Institute, Republic of Korea.

## FIGURE LEGENDS

## REFERENCES

1. Addison M, Xu Q, Cayuso J, Wilkinson DG. 2018. Cell Identity Switching Regulated by Retinoic Acid Signaling Maintains Homogeneous Segments in the Hindbrain. Dev Cell 45: 606–620 e603.

2. Arda O, Göksügür N, Tüzün YJCid. 2014. Basic histological structure and functions of facial skin. 32: 3–13.

3. Austin E, Koo E, Merleev A, Torre D, Marusina A, Luxardi G, Mamalis A, Isseroff RR, Ma’ayan A, Maverakis E et al. 2021. Transcriptome analysis of human dermal fibroblasts following red light phototherapy. Sci Rep 11: 7315.

4. Awgulewitsch A. 2003. Hox in hair growth and development. Naturwissenschaften 90: 193–211.

5. Baker DJ, Wijshake T, Tchkonia T, LeBrasseur NK, Childs BG, Van De Sluis B, Kirkland JL, Van Deursen JMJN. 2011. Clearance of p16Ink4a-positive senescent cells delays ageing-associated disorders. 479: 232–236.

6. Baumann L, Bernstein EF, Weiss AS, Bates D, Humphrey S, Silberberg M, Daniels R. 2021. Clinical Relevance of Elastin in the Structure and Function of Skin. Aesthet Surg J Open Forum 3: ojab019.

7. Brown TM, Krishnamurthy K. 2021. Histology, dermis. In StatPearls [Internet]. StatPearls Publishing.

8. Casella G, Munk R, Kim KM, Piao Y, De S, Abdelmohsen K, Gorospe M. 2019. Transcriptome signature of cellular senescence. Nucleic Acids Res 47: 11476.

9. Chung CL, Lawrence I, Hoffman M, Elgindi D, Nadhan K, Potnis M, Jin A, Sershon C, Binnebose R, Lorenzini AJG. 2019. Topical rapamycin reduces markers of senescence and aging in human skin: an exploratory, prospective, randomized trial. 41: 861–869.

10. Contet-Audonneau J, Jeanmaire C, Pauly GJBJoD. 1999. A histological study of human wrinkle structures: comparison between sun-exposed areas of the face, with or without wrinkles, and sun-protected areas. 140: 1038–1047.

11. de Farias Pires T, Azambuja AP, Horimoto ARVR, Nakamura MS, de Oliveira Alvim R, Krieger JE, Pereira ACJC, Cosmetic, Dermatology I. 2016. A population-based study of the stratum corneum moisture. 79–87.

12. Elias PM, Choi EHJEd. 2005. Interactions among stratum corneum defensive functions. 14: 719–726.

13. Ezure T, Sugahara M, Amano SJB. 2019. Senescent dermal fibroblasts negatively influence fibroblast extracellular matrix-related gene expression partly via secretion of complement factor D. 45: 556–562.

14. Farage MA, Miller KW, Elsner P, Maibach HIJC, toxicology o. 2007. Structural characteristics of the aging skin: a review. 26: 343–357.

15. Garner WLJP, surgery r. 1998. Epidermal regulation of dermal fibroblast activity. 102: 135–139.

16. Ghadially R, Brown BE, Sequeira-Martin SM, Feingold KR, Elias PMJTJoci. 1995. The aged epidermal permeability barrier. Structural, functional, and lipid biochemical abnormalities in humans and a senescent murine model. 95: 2281–2290.

17. Haeusler HJSJ. 2015. Efficacy of a hyaluronic acid gel to improve the skin properties. 141: 16–18.

18. Hamada Ji, Omatsu T, Okada F, Furuuchi K, Okubo Y, Takahashi Y, Tada M, Miyazaki YJ, Taniguchi Y, Shirato HJIjoc. 2001. Overexpression of homeobox gene HOXD3 induces coordinate expression of metastasis-related genes in human lung cancer cells. 93: 516–525.

19. Heinz AJArr. 2021. Elastic fibers during aging and disease. 66: 101255.

20. Holland PWJWIRDB. 2013. Evolution of homeobox genes. 2: 31–45.

21. Kim M, Kim SM, Kwon S, Park TJ, Kang HYJED. 2019. Senescent fibroblasts in melasma pathophysiology. 28: 719–722.

22. Kömüves LG, Ma XK, Stelnicki E, Rozenfeld S, Oda Y, Largman CJDdaopotAAoA. 2003. HOXB13 homeodomain protein is cytoplasmic throughout fetal skin development. 227: 192–202.

23. Krutmann J, Schikowski T, Morita A, Berneburg MJJoID. 2021. Environmentally-induced (extrinsic) skin aging: Exposomal factors and underlying mechanisms. 141: 1096–1103.

24. La Celle PT, Polakowska RRJJoBC. 2001. Human Homeobox HOXA7 Regulates KeratinocyteTransglutaminase Type 1 and Inhibits Differentiation. 276: 32844–32853.

25. Lago JC, Puzzi MBJPO. 2019. The effect of aging in primary human dermal fibroblasts. 14: e0219165.

26. Lavker RM, Zheng P, Dong GJCiGM. 1989. Morphology of aged skin. 5: 53–67.

27. Lee YI, Choi S, Roh WS, Lee JH, Kim T-GJIJoMS. 2021. Cellular senescence and inflammaging in the skin microenvironment. 22: 3849.

28. Liu QN, Tang YY, Zhao JR, Li YT, Yang RP, Zhang DZ, Cheng YX, Tang BP, Ding F. 2021. Transcriptome analysis reveals antioxidant defense mechanisms in the red swamp crayfish Procambarus clarkia after exposure to chromium. Ecotoxicol Environ Saf 227: 112911.

29. Mack JA, Abramson SR, Ben Y, Coffin JC, Rothrock JK, Maytin EV, Hascall VC, Largman C, Stelnicki EJJTFJ. 2003. Hoxb13 knockout adult skin exhibits high levels of hyaluronan and enhanced wound healing. 17: 1352–1354.

30. Mallo M, Wellik DM, Deschamps J. 2010. Hox genes and regional patterning of the vertebrate body plan. Dev Biol 344: 7–15.

31. Morgan RJTig. 2006. Hox genes: a continuation of embryonic patterning? 22: 67–69.

32. Pearson JC, Lemons D, McGinnis WJNRG. 2005. Modulating Hox gene functions during animal body patterning. 6: 893–904.

33. Pfisterer K, Shaw LE, Symmank D, Weninger W. 2021. The Extracellular Matrix in Skin Inflammation and Infection. Front Cell Dev Biol 9: 682414.

34. Proksch E, Brandner JM, Jensen JMJEd. 2008. The skin: an indispensable barrier. 17: 1063–1072.

35. Quinonez SC, Innis JWJMg, metabolism. 2014. Human HOX gene disorders. 111: 4–15.

36. Ressler S, Bartkova J, Niederegger H, Bartek J, Scharffetter-Kochanek K, Jansen-Dürr P, Wlaschek MJAc. 2006. p16INK4A is a robust in vivo biomarker of cellular aging in human skin. 5: 379–389.

37. Rübe CE, Bäumert C, Schuler N, Isermann A, Schmal Z, Glanemann M, Mann C, Scherthan HJnA, Disease Mo. 2021. Human skin aging is associated with increased expression of the histone variant H2A. J in the epidermis. 7: 7.

38. Rux DR, Wellik DMJDD. 2017. Hox genes in the adult skeleton: Novel functions beyond embryonic development. 246: 310–317.

39. Serpente P, Tumpel S, Ghyselinck NB, Niederreither K, Wiedemann LM, Dolle P, Chambon P, Krumlauf R, Gould AP. 2005. Direct crossregulation between retinoic acid receptor beta and Hox genes during hindbrain segmentation. Development 132: 503–513.

40. Shehwana H, Ijaz S, Fatima A, Walton S, Sheikh ZI, Haider W, Naz S. 2021. Transcriptome Analysis of Host Inflammatory Responses to the Ectoparasitic Mite Sarcoptes scabiei var. hominis. Front Immunol 12: 778840.

41. Shuster S, Black MM, McVitie EJBJoD. 1975. The influence of age and sex on skin thickness, skin collagen and density. 93: 639–643.

42. Stelnicki EJ, Arbeit J, Cass DL, Saner C, Harrison M, Largman CJJoid. 1998. Modulation of the human homeobox genes PRX-2 and HOXB13 in scarless fetal wounds. 111: 57–63.

43. Taszkun I, Tomaszewska E, Dobrowolski P, Zmuda A, Sitkowski W, Muszynski S. 2019. Evaluation of Collagen and Elastin Content in Skin of Multiparous Minks Receiving Feed Contaminated with Deoxynivalenol (DON, vomitoxin) with or without Bentonite Supplementation. Animals-Basel 9.

44. Tobin DJJJotv. 2017. Introduction to skin aging. 26: 37–46.

45. Tzaphlidou MJM. 2004. The role of collagen and elastin in aged skin: an image processing approach. 35: 173–177.

46. Varani J, Dame MK, Rittie L, Fligiel SE, Kang S, Fisher GJ, Voorhees JJJTAjop. 2006. Decreased collagen production in chronologically aged skin: roles of age-dependent alteration in fibroblast function and defective mechanical stimulation. 168: 1861–1868.

47. Victorelli S, Lagnado A, Halim J, Moore W, Talbot D, Barrett K, Chapman J, Birch J, Ogrodnik M, Meves AJTEj. 2019. Senescent human melanocytes drive skin ageing via paracrine telomere dysfunction. 38: e101982.

48. Waaijer ME, Gunn DA, Adams PD, Pawlikowski JS, Griffiths CE, Van Heemst D, Slagboom PE, Westendorp RG, Maier ABJJoGSABS, Sciences M. 2016. P16INK4a positive cells in human skin are indicative of local elastic fiber morphology, facial wrinkling, and perceived age. 71: 1022–1028.

49. Waaijer ME, Parish WE, Strongitharm BH, van Heemst D, Slagboom PE, de Craen AJ, Sedivy JM, Westendorp RG, Gunn DA, Maier ABJAc. 2012. The number of p16INK4a positive cells in human skin reflects biological age. 11: 722–725.

50. Waller JM, Maibach HIJSr, technology. 2005. Age and skin structure and function, a quantitative approach (I): blood flow, pH, thickness, and ultrasound echogenicity. 11: 221–235.

51. Wang AS, Dreesen OJFiG. 2018. Biomarkers of cellular senescence and skin aging. 9: 247.

52. Wang Z, Man M-Q, Li T, Elias PM, Mauro TMJA. 2020. Aging-associated alterations in epidermal function and their clinical significance. 12: 5551.

53. Wu JH, Yan ZW, Husile, Zhang WG, Li JQ. 2010. [Hoxc13 and the development of hair follicle]. Yi Chuan 32: 656–662.

54. Yu Q, Kilik U, Holloway EM, Tsai YH, Harmel C, Wu A, Wu JH, Czerwinski M, Childs CJ, He Z et al. 2021. Charting human development using a multi-endodermal organ atlas and organoid models. Cell 184: 3281–3298 e3222.

55. Zhang Y, Wang Y, Li Y, Yang Y, Jin M, Lin X, Zhuang Z, Guo K, Zhang T, Tan WJG. 2023. Application of Collagen-Based Hydrogel in Skin Wound Healing. 9: 185.

